# Rhythmic replay of short-term memory neural patterns revealed by time-resolved error prediction

**DOI:** 10.64898/2026.06.22.733876

**Authors:** Nikolay Syrov, Skyla Schmidt, Robin Rademacher, Xenia Kobeleva

## Abstract

Theta oscillations are hypothesized to provide a temporal scaffold for short-term memory (STM). In this model, memory representations are organized into successive theta phases, reducing conflict between competing representations during encoding and maintenance. Previous studies have shown that sensory representations of memorized information are rhythmically reactivated at theta frequency. Whether this theta-rhythmic mechanism is shared across encoding and maintenance, traditionally viewed as temporally distinct processes, remains unclear. It is also unknown whether this rhythmic reactivation predicts subsequent memory fidelity rather than merely reflecting the reinstatement of sensory representations. We recorded EEG from 21 participants performing STM with colored, oriented items under low and high memory load. Using time-resolved multivariate pattern analysis, we predicted subsequent STM error from encoding- and maintenance-period activity. We found that distributed neural patterns predicted STM error and that this prediction fluctuated at theta frequency, independently of memory load and feature. Cross-temporal generalization indicated that a shared neural pattern recurred across encoding and into maintenance, consistent with periodic reactivation of a common neural code rather than a succession of distinct states. Prediction was temporally offset across spatial positions and item features, indicating that the features of a multi-feature object are encoded by separate, rhythmically interleaved processes. These findings characterize STM encoding and maintenance as a rhythmic, recurrent process and link theta-rhythmic fluctuations in memory fidelity to distributed cortical activity spanning multiple brain regions and frequency bands.

## Introduction

Short-term memory (STM) is the capacity to temporarily store a limited amount of information for ongoing cognitive purposes. One long-standing question is how the brain coordinates the encoding and maintenance of multiple items, each defined by multiple features, into stable yet separable memory traces.

Multi-frequency neural oscillations have been established as neural correlates of STM, particularly theta-band activity and higher-frequency oscillations in the beta and gamma ranges (Lisman & Idiart, 1995; Lundqvist et al., 2016). Theta power has been repeatedly associated with successful encoding, with stronger theta activity for subsequently remembered than forgotten items (Hanslmayr & Staudigl, 2014; Sederberg et al., 2003). Delay-period theta increases have further been linked to the maintenance of task-relevant information (Hsieh & Ranganath, 2014; Jensen & Tesche, 2002). Neurophysiologically, theta is hypothesized to coordinate communication across distributed cortical and hippocampal regions and to govern the timing of faster oscillations, such that memory function relies not on a single locus but on a distributed network with multi-frequency interactions (Sauseng et al., 2010; Staresina & Wimber, 2019).

Beyond the mechanistic, spatiotemporal organization of memory network interactions, theta has been proposed to organize how mnemonic information is structured in time - how successive items are coded, how concurrent representations are kept apart while being continuously replayed during maintenance (Abdalaziz et al., 2023; Lisman & Jensen, 2013). While serving different functions, encoding and maintenance are not cleanly separated successive stages: maintenance begins as material is encoded, and theta-phase organization of individual items is already evident during encoding itself (Han et al., 2026). On this view, the encoding of a multi-item scene and the replay of its encoded representation during maintenance are interleaved within an ongoing theta rhythm, rather than separated into discrete epochs.

Direct evidence for theta-phase modulation of the interaction between encoding and replay of short-term memory representations in humans remains limited. Previous EEG studies show that the subjectively determined moment of successful recollection is locked to a particular theta phase (Abdalaziz et al., 2023; Ter Wal et al., 2021). Furthermore, time-resolved MVPA of M-/EEG has indicated that memory content can be rhythmically decoded from distributed cortical patterns: classification accuracy for stored content -- for instance, indoor versus outdoor scenes, or animate versus inanimate objects -- fluctuates at theta frequency and peaks at a specific phase of frontal and hippocampal theta (Fuentemilla et al., 2010; Kerrén et al., 2018).

However, three questions have remained unanswered by previous studies. First, it is unknown whether decoding fluctuations track the fidelity of memory or merely reflect the periodic reinstatement of sensory content, irrespective of recall accuracy. Second, it is unclear whether the fluctuations index a sequential sweep through the visual scene -- i.e., successive, distinct neural patterns processing different content -- or the genuine recurrence of a single reinstated pattern. Distinguishing these requires testing whether the recurring code is the same across time, which cross-temporal generalization can address (King & Dehaene, 2014). Third, because previous MVPA studies classified whole scenes, it remains unknown whether one theta-rhythmic process governs an entire multi-item array or whether separable rhythmic dynamics can be resolved for individual items and for the distinct features within each item.

In the present study, we addressed these questions by applying time-resolved MVPA to predict continuous memory error directly from encoding and maintenance period EEG. Participants encoded arrays of items defined by color and orientation under two memory loads (two or four spatially distanced items). After a delay, one of these was cued and participants had to report its features on a continuous scale. Rather than classifying which stimulus was held in memory, we trained predictors to estimate the magnitude of subsequent recall error for the color and orientation features of cued items, and asked when, and through which oscillatory activity, the encoding and maintenance pattern carried this information.

This shift from decoding content to predicting fidelity speaks to the first question, i.e., whether the rhythmic organisation of STM is associated with recall accuracy. To address the second question -- whether recurring prediction reflects a sequential sweep of distinct neural codes or the recurrence of a single code -- we applied cross-temporal generalization, testing whether a predictor trained at one moment of encoding generalizes to later moments including maintenance. To address the third, we resolved prediction separately for each item and for its two features (color and orientation), allowing us to ask whether one rhythmic process governs the whole array or whether separable dynamics operate at the level of individual items and features.

Together, our study contributes to a better understanding of the rhythmic, recurrent organization of multi-item STM. As such, it has implications for computational models of STM and temporally targeted precise neuromodulation of memory processes.

## Methods

### Key resources table

| RESOURCE | SOURCE | IDENTIFIER |
| --- | --- | --- |
| **Software and algorithms** |  |  |
| MNE-Python | Gramfort et al., 2013 | https://mne.tools/stable/install/index.html |
| Autoreject | Jas et al., 2017 | https://github.com/autoreject/autoreject |
| Working memory mixture model (three-component) | Bays et al., 2009 | https://github.com/smathot/biased_memory_toolbox |
| PsychoPy | Peirce et al., 2019 | https://psychopy.org/ |
| Preprocessing and analysis scripts | This paper | [GitHub URL will be publicly available as of the date of publication] |

### Participants

Twenty-one participants (11 female; 10 male, mean age 25, range 19–34) took part in this study. All, except one participant, were right-handed, English speaking and had normal or corrected-to-normal vision. The study was approved by the Ruhr University Bochum Research Ethics Committee (Protocol number: 2025-416-f-S). All participants provided written informed consent prior to participation and were informed about the experimental procedure.

### Short-term memory task design

Task and stimuli. Participants performed a delayed continuous-report visual STM task, which we implemented in PsychoPy (Peirce, 2007). Each trial began with a fixation cross of jittered duration (800–1300 ms), followed by the encoding period, during which the encoding array was presented. The array consisted of either 2 (low load) or 4 (high load) colored, oriented rectangles, each presented at one of five fixed locations (upper-left, upper-right, lower-left, lower-right, and center). The occupied locations varied randomly from trial to trial, so that participants could not anticipate at which positions items would appear. For each trial, item colors and orientations were constrained to be mutually distinct. Colors were sampled from a circular CIELAB hue space (L = 60, chroma = 38) with a minimum pairwise separation of ∼79°. Orientations were selected from a discrete set ranging from 0° to 170° in 10° increments, with a minimum pairwise separation of 30°.

The encoding period lasted 800 ms and was followed by a 700 ms maintenance interval, a 300 ms visual mask, and a subsequent 1000 ms delay.

After the delay, a spatial retro-cue was presented for 2000 ms, indicating which of the encoded items was to be reported. Because the cued item was unpredictable, participants were required to maintain all items. Following the cue, participants reported the remembered features of the cued item using continuous responses: color was selected on a continuous color wheel with the computer mouse, and orientation was reproduced by rotating a rectangle using the mouse wheel. Response time was unrestricted. The order of color and orientation reports was randomized across trials, minimizing potential order effects on memory fidelity.

Design and procedure. Each block contained 40 trials, with 10-second breaks after every 10 trials. The session comprised four blocks: three high-load (4 items) and one low-load (2 items). This resulted in a total of 160 trials (120 high-load and 40 low-load). The order of the blocks was randomized between subjects. Before the main task, participants received instructions, were asked to minimize blinking during the encoding and maintenance phase, and completed a 5-trial practice run to confirm task understanding. Including EEG cap preparation, each session lasted no more than 2 hours.

**Figure 1.**
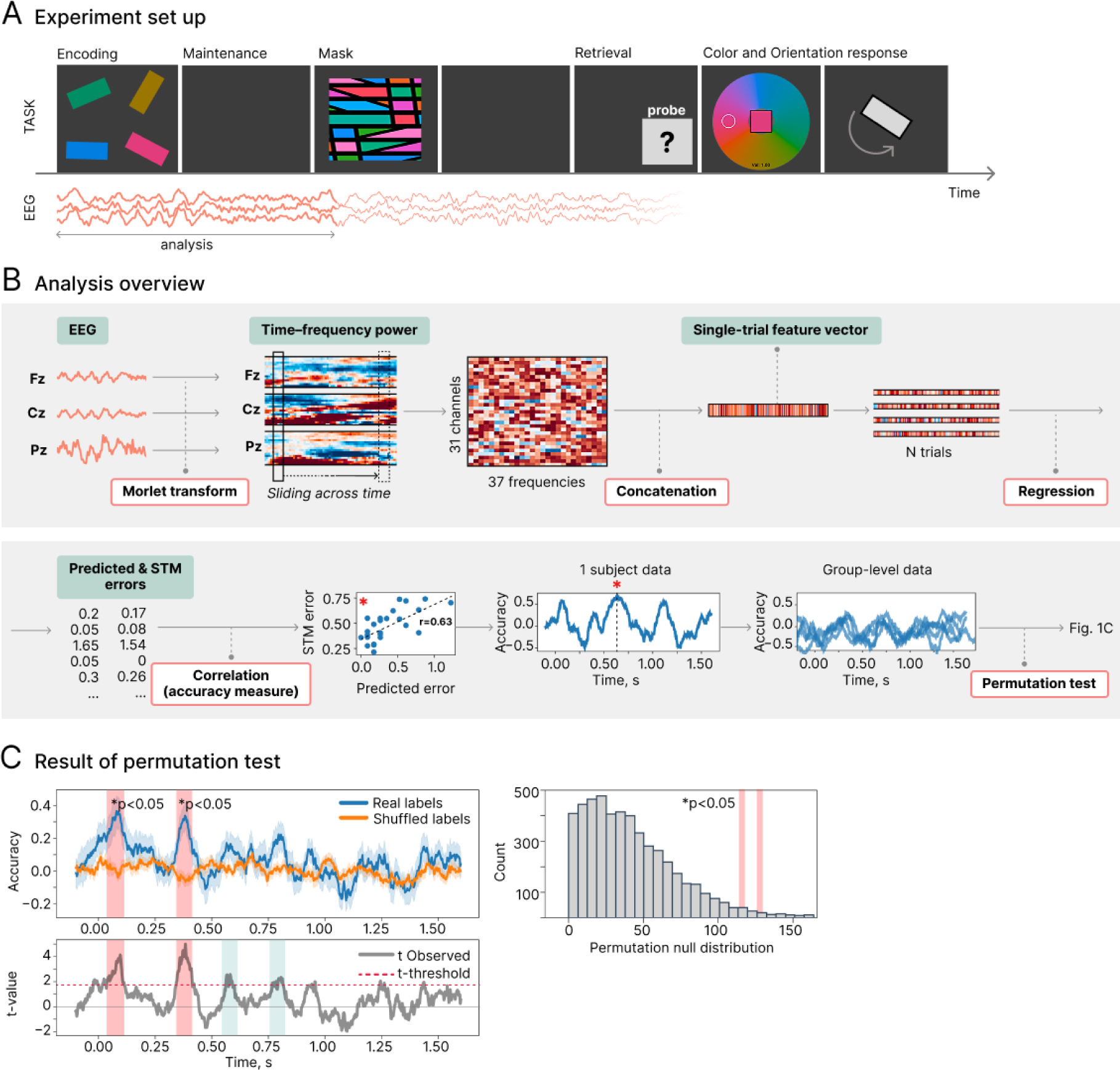
Delayed continuous-report visual short-term memory task and multivariate decoding analysis pipeline. **A.** Delayed continuous-report STM task. Participants encoded two or four colored, oriented rectangles presented at fixed spatial locations, maintained them across a delay, and reported the color and orientation of a retrospectively cued item using continuous-response measures. **B.** Time-resolved decoding analysis. EEG signals with evoked activity subtracted were transformed into time–frequency power using Morlet wavelets. At each time point, channel × frequency power values were concatenated into single-trial feature vectors and entered into a ridge-regression model to predict trial-by-trial memory error separately for color and orientation. Model accuracy was quantified as the correlation between predicted and observed STM errors. The red asterisk indicates an example prediction peak, with the corresponding scatterplot showing the correlation between observed memory errors and errors predicted from the neural pattern at that time point. **C.** Group-level statistical analysis. Decoding accuracy was quantified as the correlation between observed and predicted errors and evaluated over time. Statistical significance was assessed using a one-sided cluster-based permutation test, comparing observed decoding performance against a surrogate distribution obtained from shuffled error labels.

### Behavioral data analysis

Color responses and targets were expressed in radians on a circular hue wheel (full wheel = 2π), whereas orientation responses and targets were expressed in radians within a 0–π range. Color errors were computed as signed circular distances wrapped to (−π, π], and orientation errors as signed minimal distances in 180°-periodic space, wrapped to (−π/2, π/2]. Absolute errors were defined as the magnitudes of these signed errors.

To assess dependence between feature-specific errors, for each participant, mutual information (MI) between absolute color and orientation errors was estimated using a k-nearest-neighbor estimator. The group statistic was defined as the mean MI across participants. Significance was assessed against a permutation null distribution generated by shuffling errors within each participant (1,000 permutations) and recomputing the group mean. Analyses were performed separately for each memory load.

To test whether recall error varied with target location, feature, and memory load, absolute errors were first normalized by the maximum possible error for each feature, accounting for their different circular ranges (0–π for color; 0–π/2 for orientation). Normalized absolute errors were averaged within participant, feature, and location and entered into a repeated-measures ANOVA with Feature (color, orientation), Location (five levels), and Memory load (2-, 4-items) as within-participant factors. Greenhouse–Geisser correction was applied to factors with more than two levels.

#### Mixture-model decomposition

To estimate the sources of memory error, we fit the mixture model of (Bays et al., 2009), decomposing continuous-report errors into target, swap (misbinding), and guess components, separately to color and orientation responses using the biased_memory_toolbox (Zhou et al., 2022). Participant-specific parameters were then used to derive trial-wise posterior probabilities of target, swap, and guess responses (P_hit, P_swap, P_guess) via Bayes’ rule.

### EEG recording

Participants were seated comfortably in front of a computer screen and completed practice trials to familiarize themselves with the task. EEG was recorded using an actiChamp amplifier (Brain Products, Germany) at a sampling rate of 1 kHz from 31 scalp electrodes positioned according to 10–10 system at Fp1, Fp2, F7, F3, F1, Fz, F4, F8, FC5, FC3, FC1, FCz, FC2, FC4, FC6, C3, Cz, C4, T7, T8, CP5, CPz, CP6, P7, P3, Pz, P4, P8, O1, and O2. Channel TP10 was used as the recording reference, and the data were subsequently re-referenced to the common average. Electrode impedances were maintained below 10 kΩ for all channels. EEG acquisition was synchronized with stimulus presentation using an analog photosensor attached to the screen and connected to a Micro1404 interface (Cambridge Electronic Design, UK), which was in turn linked to the EEG amplifier.

### EEG preprocessing

Preprocessing was performed in MNE-Python (Gramfort et al., 2013). Continuous EEG data were first notch-filtered at 50 Hz and its harmonics (100, 150, and 200 Hz) to suppress line noise and subsequently band-pass filtered between 0.1 and 70 Hz using a zero-phase FIR filter. Noisy channels were identified by visual inspection and repaired using spherical spline interpolation (mean = 0.38 channels per participant), after which the data were re-referenced to the common average. Independent component analysis (FastICA) was then performed to remove ocular and muscle artifacts. To improve component decomposition, ICA was fitted to a copy of the data band-pass filtered between 1 and 40 Hz and retaining 99% of variance, and the resulting decomposition was subsequently applied to the broadband (0.1–70 Hz) data. Components corresponding to eye-movement and muscle artifacts were identified by visual inspection and removed.

Continuous data were segmented into epochs from −1 to +2 s relative to memory-array onset, covering the encoding and maintenance intervals. Epochs were baseline-corrected using the −200 to 0 ms pre-stimulus window and downsampled to 500 Hz.

Bad epochs were detected and corrected using AutoReject (Jas et al., 2017), which repairs or rejects epochs based on cross-validated, channel-specific peak-to-peak amplitude thresholds (consensus κ and the number of interpolated channels ρ optimized by cross-validation over κ ∈ [0.5, 1.0] and ρ ∈ {0, 1, 2, 3}). Channels that exceeded their threshold within an epoch were flagged. Epochs were rejected when the proportion of flagged channels exceeded the consensus criterion (κ), and otherwise retained with up to three affected channels interpolated. On average, 2 epochs (1.25%) were rejected per participant.

### Time-Frequency Decomposition and Induced Spectral Power

Induced (non-phase-locked) spectral power reflecting oscillatory activity without contribution of evoked response was obtained (Luu et al., 2004; Trujillo & Allen, 2007). This was done with a single-trial subtraction approach via removing the evoked response directly from each trial’s complex spectral representation prior to power computation (Kalcher & Pfurtscheller, 1995). Time-frequency spectral power dynamics were then performed using complex Morlet wavelet transform. The frequencies of the wavelets ranged from 3 Hz to 40 Hz with 1 Hz step. The full-width at half-maximum (FWHM) was 187 ms.

### Time-resolved error prediction

Time-resolved multivariate pattern analysis (MVPA) was used to predict trial-wise recall error from the spatial–spectral distribution of oscillatory power. Decoding was performed separately for color and orientation errors and, within each feature dimension, separately for each cued spatial position. Because two or four items were presented simultaneously at fixed locations and only one item was retrospectively cued for report, position-specific decoding was motivated by the hypothesis that items at different locations are sampled with distinct temporal dynamics during encoding.

For each participant and time point, feature vectors were constructed by concatenating power values across channels and frequencies. Features were standardized within each cross-validation fold and entered into an L2-regularized linear regression model (ridge regression; α = 1.0) trained to predict absolute STM error, defined as the magnitude of the wrapped circular distance between the reported and target feature value (0–π for color; 0–π/2 for orientation). Model performance was evaluated using 5-fold cross-validation. Decoding accuracy was assessed as the Pearson correlation between predicted and observed errors across folds. Repeating this analysis at successive time points (step size = 1 sample) yielded a time-resolved decoding-accuracy time course. Surrogate accuracy time courses were generated by randomly permuting error values across trials and repeating the full 5-fold cross-validated MVPA procedure. Five independent label permutations were performed per participant, spatial position, and feature dimension. Each permutation was evaluated using cross-validation, and surrogate performance was averaged across all folds and permutations.

#### Cross-subject generalization

To test whether neural patterns predictive of recall error generalized across individuals, we used subject-wise cross-validation. At each time point, a ridge regression model was trained on data from all but one participant and used to predict the absolute recall error of the held-out participant, yielding one prediction time course per participant. Surrogate time courses were generated using the same error-shuffling procedure as in the within-subject analysis.

Group-level statistics. Group-level significance was assessed using a one-sided cluster-based permutation test across time (5,000 permutations). Clusters were defined using a cluster-forming threshold t = 1.725 (p < 0.05, tail = 1; df = 20) and considered significant at the cluster level when p < 0.05.

#### Feature contribution analysis

To characterize which channels and frequencies were most contributing in color and orientation error prediction, we examined the coefficients used for MVPA models. For each participant, regression weights were extracted from the time windows showing significant group-level decoding. Within each model, weights were z-scored across features and averaged across the selected time windows and across target item positions (color and orientation were analyzed separately to test if there are feature-specific patterns). The resulting per-participant channel × frequency maps were tested against zero at the group level with a two-sided cluster-based permutation test (10000 permutations; cluster-forming threshold t = 2.0; MNE-Python spatio_temporal_cluster_1samp_test), using channel adjacency obtained from the montage via mne.channels.find_ch_adjacency so that clusters could extend across neighboring channels and adjacent frequency bins.

#### Cross-temporal generalization of error prediction

To assess the temporal stability of the neural patterns predicting recall error, we performed a cross-temporal generalization (CTG) analysis (King & Dehaene, 2014). Following the same MVPA pipeline and cross-validation procedure as the main time-resolved decoding analysis, a ridge model trained at each encoding time point was tested at every other time point, which resulted in a train × test generalization matrix, where each cell contains the Pearson correlation between predicted and observed error. Group-level matrices were computed separately for color and orientation and for each target spatial position and tested against zero with a one-sided cluster-based permutation test (10,000 permutations; tail = 1; cluster-forming threshold t = 1.725, p < .05 one-sided, df = 20). Significant prediction confined to the diagonal (train time = test time) would indicate a succession of transient, temporally specific encoding codes, whereas off-diagonal clusters separated from the diagonal would indicate reactivation of an earlier code; a broad, sustained off-diagonal (square) pattern would indicate a stable maintained code.

### Temporal relationships between STM error predictions across spatial locations and features

Phase differences across spatial locations. To test whether the 4 Hz phase of the decoding rhythm varied with target location, a 4 Hz sinusoid was fitted to each participant’s decoding time course separately for each of the five target coordinates, and the phase of the fitted oscillation was extracted. To remove inter-individual differences in absolute phase, each participant’s coordinate phases were expressed relative to that participant’s circular mean phase across coordinates. The centered phases were entered into a one-way Watson–Williams circular ANOVA with target coordinate as a five-level factor.

Color and orientation error prediction phase coupling analysis. To test whether the rhythmic fluctuations in color and orientation decoding were temporally coordinated, we assessed whether the two features maintained a consistent phase offset across participants, pooling spatial coordinates as repeated observations. Each time course was linearly detrended (scipy.signal.detrend), mean-subtracted, and multiplied by a Tukey window (α = 0.5). We extracted the 4 Hz phase of each decoding time course from its complex Fourier coefficient at 4 Hz, and took the color–orientation phase difference (Δφ = φ_color − φ_orientation) for each participant and coordinate.

Because different target item spatial coordinates within a participant are not independent, group-level consistency of Δφ was tested with a subject-blocked permutation Rayleigh test (Rayleigh test for non-uniformity; (Berens, 2009): the observed Rayleigh statistic over all subject–coordinate observations was compared with a null distribution (10,000 permutations) in which each participant’s complete set of coordinate phase differences was rotated by a single random angle, preserving within-subject structure while removing group-level phase alignment. A significant result of this testing would mean Δφ clustered around a consistent direction across participants rather than being uniformly distributed — i.e., color and orientation decoding held a reliable, non-random 4 Hz phase relationship at the group level. As a complementary test of whether Δφ was organized by target location rather than reflecting a single global offset, we ran a within-subject condition-pattern permutation test. The statistic was the summed resultant length of Δφ across coordinates, compared with a null distribution (10,000 permutations) in which coordinate labels were shuffled within each participant. A significant result would indicate that the color–orientation phase offset varied systematically across spatial coordinates, whereas a non-significant result indicates a consistent offset that does not depend on target location.

## Results

### Behavioral results

Mean absolute color error was 15.7° ± 6.4° (0.27 ± 0.11 rad) in the low-load condition and 32.8° ± 18.3° (0.57 ± 0.32 rad) in the high-load condition. Mean absolute orientation error was 12.5° ± 9.7° (0.22 ± 0.17 rad) and 24.6° ± 9.5° (0.43 ± 0.17 rad) for low and high load, respectively. Response times did not differ markedly across conditions (color: 2.9 s vs. 2.6 s; orientation: 3.7 s vs. 3.5 s), with orientation reports taking consistently longer than color reports. Individual distributions of memory errors for color and orientation across participants are presented in Appendix.

Color and orientation errors showed no significant statistical dependence: mutual information did not exceed the within-participant permutation null at either low load (p = 0.43) or high load (p = 0.07). There was no significant difference in the spatial distribution of STM errors: there was no main effect of Location (F(4,80) = 0.87, p = .49) and no interaction of Location with Feature (F(4,80) = 1.31, p = .27) or Load (F(4,80) = 1.55, p = .20), indicating that memory precision was spatially uniform across the screen. As expected, memory error increased with memory load (F(1,20) = 93.9, p < .001), and was lower for color than orientation (F(1,20) = 10.08, p = .005), with this feature advantage larger at higher load (Feature × Load: F(1,20) = 6.32, p = .021).

**Figure 2.**
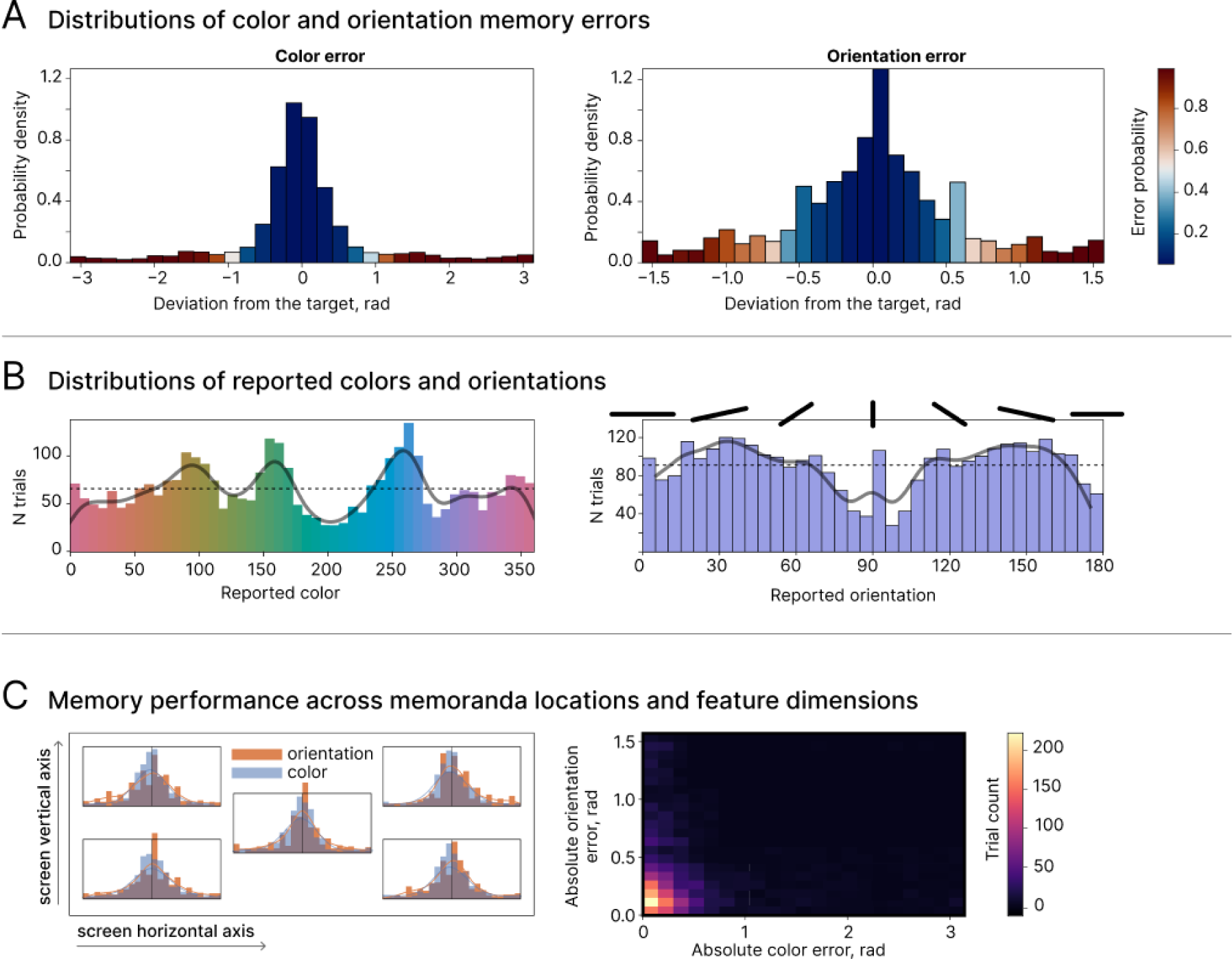
Behavioral STM performance across sensory features and memoranda locations. **A.** Distributions of color and orientation memory errors. Bars are color-coded according to the fitted model-based error probability (EP), with higher values indicating a greater probability that the response reflected an error (forgotten) item. Orientation errors were more broadly distributed than color errors. **B.** Group-level distributions of reported colors and orientations. The non-uniform structure of both distributions suggests a preference of color and orientation values within each category. **C.** Memory errors control analyses. Left: color and orientation error distributions are shown separately for different spatial positions. No reliable differences in error magnitude were observed across positions. Right: two-dimensional distribution of absolute color and orientation errors.

### Time-resolved error prediction results

Time-resolved MVPA identified multiple temporal windows within the encoding period, in which neural patterns significantly predicted subsequent memory error. In both memory load conditions, prediction of color and orientation error exhibited an oscillating pattern. Figure 3,A presents the group-level prediction time courses with significant clusters identified by cluster-based permutation testing; a similar pattern was observed for within-subject regression models (see Appendix for details).

Peak frequency analysis revealed that memory error prediction for both color and orientation fluctuated at a mean frequency of 4 Hz (range: 3–5 Hz) for both low and high-load conditions (Figure 3,B).

**Figure 3.**
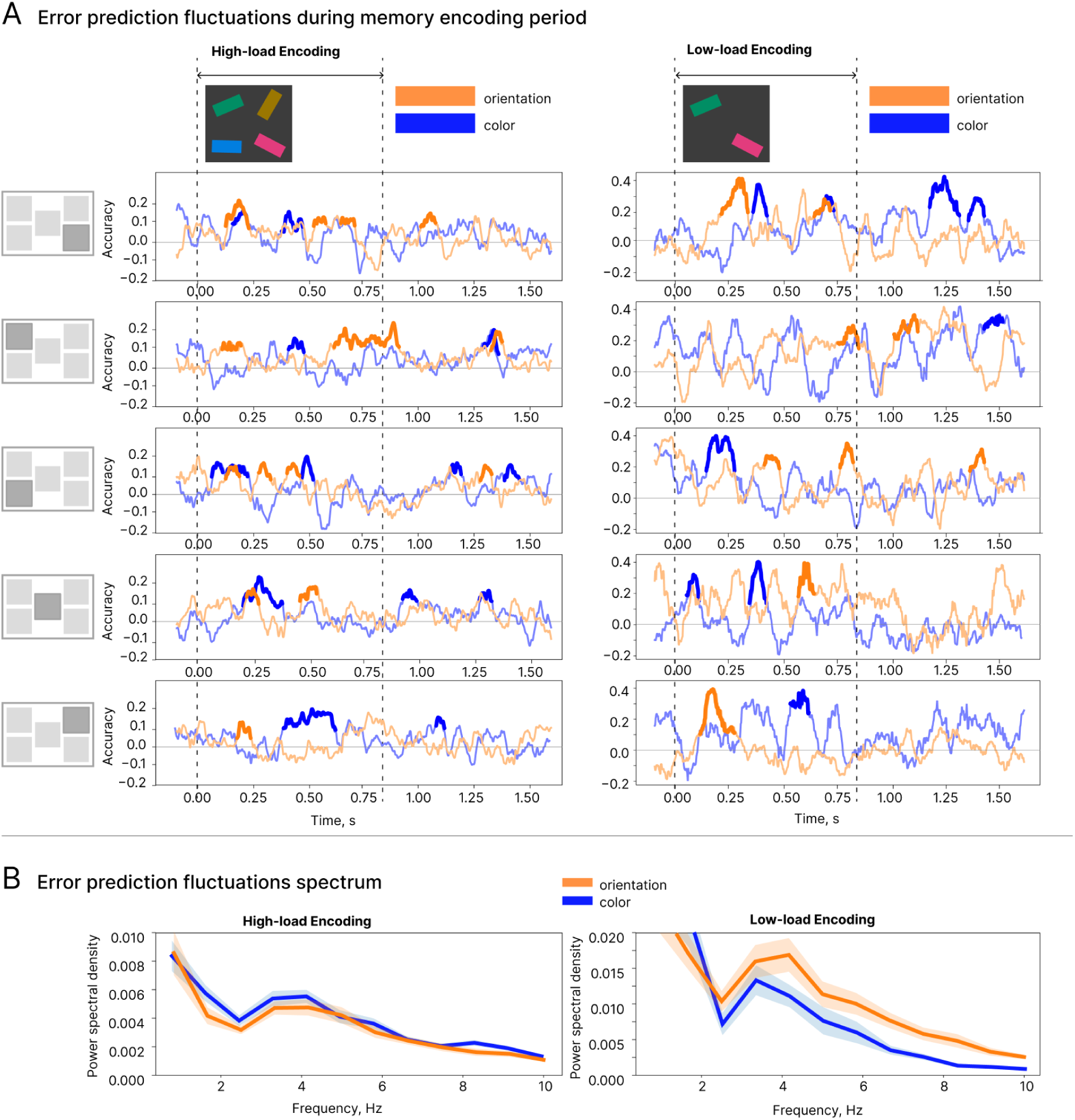
Temporal and spectral characteristics of cross-subject generalized MVPA prediction of color and orientation memory errors during the encoding period. **A.** Group-level time courses of error prediction for color (blue) and orientation (orange) in the high-load (left) and low-load (right) conditions, shown separately for different spatial positions. Prediction values are shown across the encoding period and the following maintenance interval. The dashed lines mark encoding onset and offset. In both load conditions, prediction of color and orientation errors exhibited an oscillating temporal pattern. Thickened line segments indicate significant temporal clusters identified with cluster-based permutation testing. **B.** Power spectral density of the prediction fluctuations, averaged across subjects and spatial positions. For both color and orientation prediction fluctuated predominantly within the 3–5 Hz range in both memory-load conditions.

#### Spatial and feature-specific phase structure of the 4 Hz decoding rhythm

A Watson–Williams circular ANOVA indicated that the mean 4 Hz phase of the error-prediction decoding time courses differed across the five target locations, F(4, 200) = 2.56, p = .039.

To test for a systematic phase shift between color and orientation encoding, we estimated the phase lag between color and orientation error-prediction dynamics across all spatial positions using circular statistics at 4 Hz. In the high-load condition, color–orientation phase differences were consistently oriented across participants (subject-blocked permutation Rayleigh test: z = 4.94, p = .036, r = 0.22), clustering around a mean offset of approximately −53° (Δφ = φ_color − φ_orientation; ≈37 ms at 4 Hz). This offset did not differ reliably across spatial positions (within-subject condition-pattern permutation: statistic = 1.51, p = .13), indicating a consistent, location-independent phase relationship. Because color carried the earlier phase (negative Δφ), these results indicate that color error prediction preceded orientation error prediction. No significant phase-lag effect was observed in the low-load condition.

**Figure 4.**
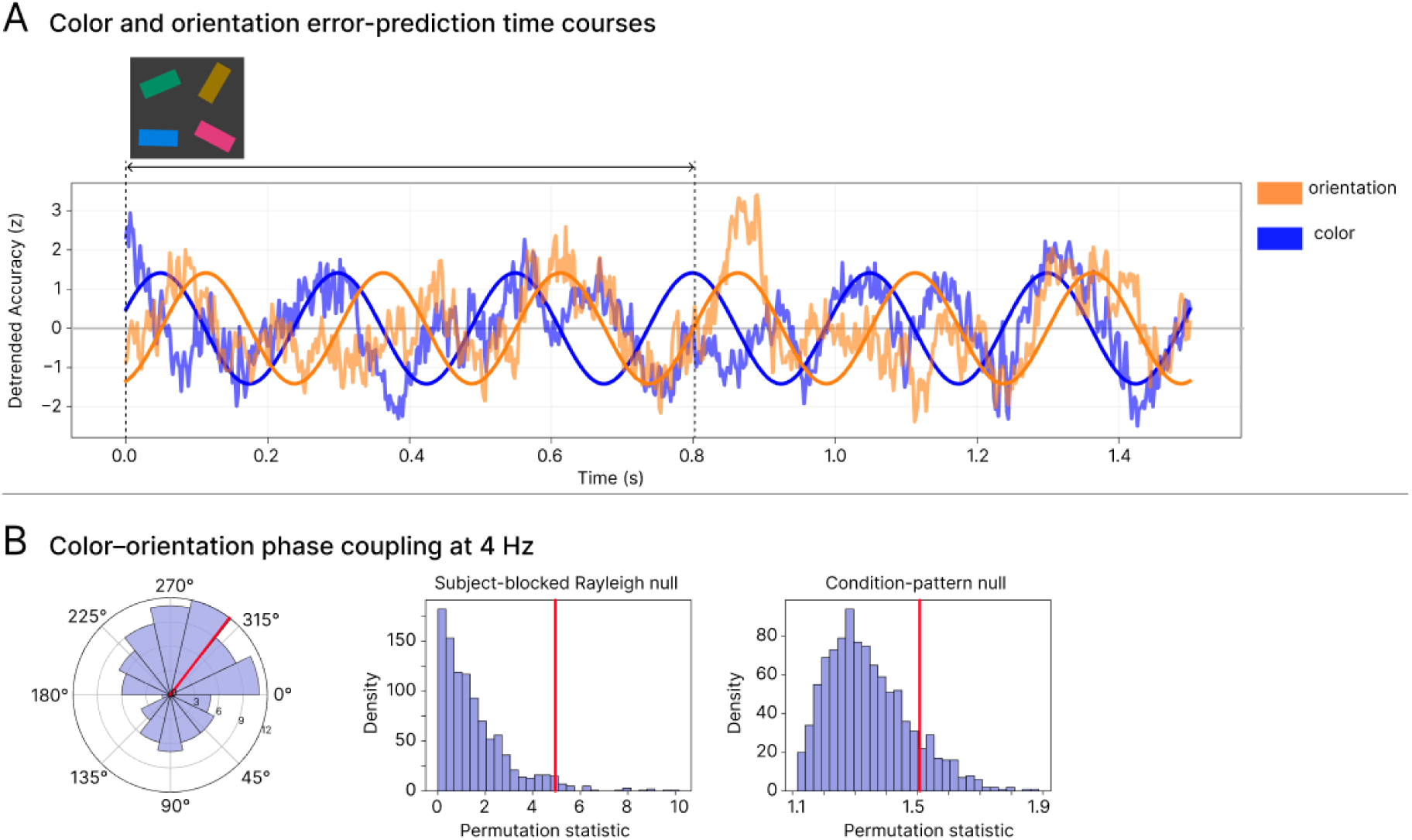
Color error prediction precedes orientation error prediction during encoding. **A.** Representative participant showing color (blue) and orientation (orange) error-prediction time courses together with fitted 4-Hz sinusoids. **B.** Color–orientation phase coupling at 4 Hz. Circular histogram of color–orientation phase differences (Äö = ö_color − φ_orientation) across participants and spatial locations; the red radial line denotes the group circular mean. Middle and right panels show permutation null distributions for the subject-blocked Rayleigh test and the within-subject condition-pattern test, respectively. Red vertical lines indicate the observed test statistics.

### Cross-temporal generalization

To determine whether the fluctuating prediction dynamics reflected reactivation of similar or distinct neural patterns, we performed a cross-temporal generalization (CTG) analysis (King & Dehaene, 2014). If the error-prediction peaks observed in the time-resolved MVPA result reflect a recurring shared neural code (i.e., similar patterns), a model trained at one prediction peak should generalize to another, producing significant off-diagonal clusters at their intersection in the time × time matrix. Conversely, if each recurrence involves a distinct pattern, significant prediction should be limited to the diagonal region.

This analysis was conducted using within-subject regression models because CTG requires neural representations to generalize across time points, making it more sensitive to temporal variability than standard decoding. Within-subject models increase sensitivity by capturing participant-specific spatial and spectral response patterns.

For both color and orientation CTG revealed significant clusters off the diagonal, indicating that a shared neural pattern predicting memory error reappears several times across the encoding and maintenance period (Figure 5, A shows CTG matrix for one spatial position, for all positions see Appendix).

**Figure 5.**
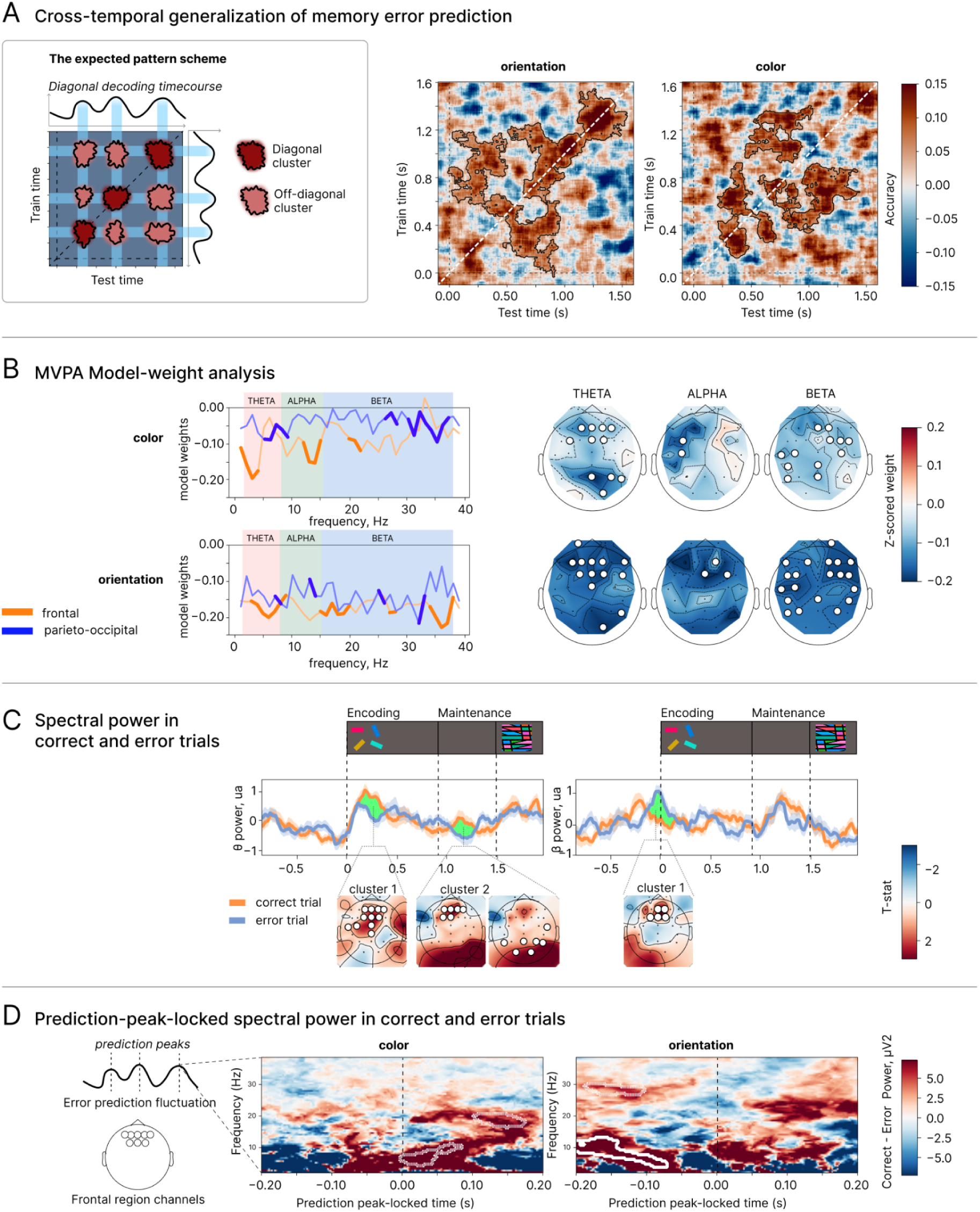
Neural representations underlying memory error prediction. Temporal generalization analysis shows periodic reinstatement of the same neural pattern. Analysis of the spatial and spectral characteristics of prediction-enabling neural patterns revealed dominant contributions of theta- and beta-band activity. **A**. Cross-temporal generalization of error-prediction decoding. *Left:* schematic of the pattern expected if a shared neural representation is periodically reinstated — a rhythmic diagonal decoding time course together with recurring off-diagonal clusters in the train-time × test-time matrix. *Right:* empirical temporal-generalization matrices for orientation and color at a single representative target coordinate show that, in addition to diagonal clusters of significant error decoding, there are also significant off-diagonal clusters. **B**. Model-weight analysis of the spectral and topographic contributions to error prediction, for color (top) and orientation (bottom). *Left:* model weights as a function of frequency for frontal (orange) and parieto-occipital (blue) channels. Bold segments indicate frequencies contributing significantly. *Right:* scalp distributions of model weights per band, with significant channels marked. **C**. Time-resolved spectral power differences between correct and error trials in the high-load condition. Correct trials showed stronger theta power during encoding and maintenance, with significant frontal and occipital clusters. Error trials showed stronger frontal beta power before stimulus onset. Statistical significance was assessed using cluster-based permutation testing. Topographical maps show significant clusters, with color indicating the corresponding t-statistic. **D**. Prediction-locked time-frequency analysis. For each participant, EEG spectral power within ±200 ms of error-prediction peaks was baseline-corrected against a surrogate distribution and contrasted between correct and error memory trials (Correct − Error power, μV^2^) using cluster-based permutation. For color error two clusters emerged at trend level, for orientation two significant clusters were found.

### Spectral and spatial contributions to STM error prediction

#### Model weight analysis

We next characterized the spatial and spectral features of error-predictive neural patterns by examining regression weights averaged across significant prediction windows. Significant contributions were distributed across theta, alpha, and beta bands and were predominantly negative, indicating that stronger oscillatory power predicted lower subsequent error (Figure 5, B). Theta-band weights showed a fronto-posterior distribution, alpha-band contributions were mainly anterior, and beta-band weights were distributed across frontal and posterior sites.

#### Spectral power analysis

To demonstrate what is happening in the raw oscillatory power in correct and error trials we first ran a pairwise comparison between spectral power dynamics during memory encoding in correct and error trials (only the high memory-load condition was analyzed, because it provided a sufficient number of memory errors for reliable comparisons).

To formally divide the trials into correct and error trials, we used a fitted mixture model, computing an error probability (EP) for each trial. Correct trials were defined as those for which the error probability for at least one feature was less than 0.25. Error trials were defined as those for which the EP for colour or orientation memory was greater than 0.75 (see Appendix for more details).

Figure 5,C shows theta power dynamics in correct and error trials during the encoding and maintenance intervals. A cluster-based permutation test revealed a significant frontal cluster during encoding, with greater theta power in correct than in error trials (p < 0.05, Cohen’s d = 1.13). During the maintenance phase, two significant clusters emerged, indicating stronger theta power in correct trials over frontal and occipital regions (p < 0.05; Cohen’s d = 1.0, 1.0, correspondingly). Additionally, in the beta frequency range a significant frontal cluster was identified. Notably, this cluster emerged before stimulus onset (p < 0.05, Cohen’s d = 1.0).

#### Prediction-locked spectral power

Additionally we compared spectral perturbations in time windows around moments when error prediction peaks. For each subject EEG spectral power in two regions of interest (frontal and parieto-occipital EEG channels) was analyzed within ±200 ms of error prediction peaks (for color and orientation separately). Peak-locked activity was baselined to surrogate signal and then compared between correct and error memory trials using the permutation cluster test. Full methodological details are provided in the Appendix.

For orientation errors, analysis of peak-locked spectral power in frontal channels identified a significant theta-band cluster (p < 0.05, Cohen’s d = 0.69), whereas an identified beta-band cluster did not reach significance (p = 0.10, Cohen’s d = 0.69). For color errors, no clusters survived correction, although trend-level theta and beta clusters were observed (theta: p = 0.09, Cohen’s d = 0.78; beta: p = 0.10, Cohen’s d = 0.79). For parieto-occipital EEG channels no clusters were identified.

## Discussion

Traditionally, encoding and maintenance have been viewed as temporally discrete processes, confined to successive post-stimulus intervals and supported by distinct brain regions and oscillatory frequencies. Prefrontal theta- and beta-oscillations have been consistently implicated in the encoding and maintenance of memory representations, and their distinct roles in these processes have been extensively discussed (Chang et al., 2023; Hanslmayr & Staudigl, 2014; Schmidt et al., 2019; Wynn et al., 2024). Recent evidence has expanded this view, suggesting that memory relies on the dynamic reactivation of distributed neural patterns through interactions across cortical networks and different oscillatory rhythms (Christophel et al., 2017; Han et al., 2026; Schmidt et al., 2019). In the present study, we provide evidence that a spatially distributed, multi-frequency neural pattern during short-term memory encoding predicts reported memory error. The present study revealed that fronto-parietal, multi-frequency neural patterns during encoding predicted subsequent memory error. Rather than being sustained or transiently appearing within a fixed time window, error prediction fluctuated at theta frequency (∼3–5 Hz), with a shared neural pattern recurring across encoding and maintenance, as shown by cross-temporal generalization. We further observed that prediction was temporally offset across spatial positions and item features, supporting prior proposals that the features of a multi-feature item are encoded separately rather than as a unified representation (Schneegans & Bays, 2017).

According to traditional theories of short-term memory, the temporal maintenance of encoded information depends on persistent neural activity established at encoding and sustained throughout the maintenance period (Sreenivasan et al., 2014). Recent findings challenge this account, demonstrating that mnemonic information can be retained in the absence of sustained firing (Kandemir et al., 2024; Rose et al., 2016; Stokes, 2015). Under activity-silent frameworks, memory traces are instead maintained within transient, spatially distributed neural circuits — for example, through short-term synaptic plasticity — that undergo theta-orchestrated reactivation during encoding and maintenance (Pals et al., 2020; Stokes, 2015). In contrast to sustained activity, such rhythmic reinstatement enables a temporal segregation of competing memory representations, thereby reducing interference in multi-item working memory (Abdalaziz et al., 2023; Fuentemilla et al., 2010; Leszczyński et al., 2015; Lisman & Jensen, 2013). Our present findings speak directly to this framework. Using time-resolved decoding of memory errors, we found that distributed, multi-frequency neural patterns associated with STM fidelity were neither sustained nor confined to a single critical moment of memory formation. Instead, these patterns recurred rhythmically at theta frequency throughout encoding and into the maintenance period.

A conceptually similar decoding approach was previously used by Fuentemilla et al. (2010), who showed that distributed occipital, frontal, and parietal activity during memory maintenance enabled differentiation of stored sensory content. Using a classification approach to predict the remembered scene category (indoor vs. outdoor), they found that classification accuracy fluctuated at theta frequency, pointing to periodic reactivation of memory content, which supports short-term retention. A comparable fluctuating structure was reported by (Kerrén et al., 2018), who found that the accuracy of time-resolved classification of neural patterns related to word-object memories fluctuated at 7–8 Hz, and was most reliable at a specific phase of the ongoing theta cycle. In the present study we used a regression approach predicting continuous memory error, and our results converge with these reports in revealing theta-rhythmic fluctuations of decoding. It is interesting to note though that Fuentemilla et al. found that the number of replay events scaled with maintenance demands, whereas we found the reinstatement dynamics to be independent of memory load. Our work further extends Fuentemilla et al. (2010) and Kerrén et al., (2018) findings through the cross-temporal generalization analysis, which revealed that the recurring prediction reflected the same neural pattern recurring cyclically throughout encoding and into maintenance, rather than a succession of distinct neural codes.

Notably, these rhythmic reinstatements have been revealed across studies using markedly different methods. **Han** et al., 2025 found that memory accuracy fluctuated with theta depending on the phase at which items were memorized, while Ter Wal et al. (2021) demonstrated that theta rhythmicity is detectable in behavioral responses themselves, with button-press timestamps indexing the moment of memory formation rhythmically modulated at 1–5 Hz. With our MVPA prediction framework, we extend this logic in a specific way: not only does the timing of memory operations fluctuate at theta, but the quality of encoding — indexed by subsequent recall error — does so as well, and we demonstrate this directly in the recurring neural patterns. In doing so, we provide direct evidence reconciling behavioral and neural oscillations. We propose that behavioral responses can serve as a proxy for the neural rhythms of memory processes.

How, then, are the different items and features of a multi-item scene or multi-feature item represented within these recurring cycles? The theta–gamma phase-coding model proposes that individual items are represented by neural assemblies firing at distinct phases of the theta cycle (Lisman & Idiart, 1995), with theta serving as a temporal scaffold within which transient high-frequency activity represents individual memory items (Jensen & Tesche, 2002; Sauseng et al., 2010; Axmacher et al., 2010; Heusser et al., 2016; Lundqvist et al., 2016; Miller et al., 2018). Consistent with the idea that theta provides a temporal scaffold for memory content, Han et al. (2026) recently reported that, in non-human primates, STM performance depended on the phase of frontal theta oscillations (3–6 Hz) at which the memory array was presented. Critically, distinct theta phases were optimal for different spatial locations, which the authors interpreted as evidence for a sequential theta wave scanning memory representations across visual space. Similarly, the theta-rhythmic STM patterns replay we observed were phase-offset across spatial locations, suggesting that spatial representations in STM may be temporally separated within the theta cycle.

Extending this spatial organization, we also found that reinstatement of STM patterns for different sensory features of the same item was temporally offset. During encoding, color information was predicted earlier than orientation information.

This temporal separation of color and orientation extends the principle of rhythmic sampling to the level of features within an item. (Fiebelkorn et al., 2013) showed that, even under sustained attention, perceptual sensitivity is rhythmically reweighted between items, with the two items sampled in antiphase at ∼4 Hz. Our findings suggest that the features of a single item are likewise sampled individually in STM, each within a separate ∼4 Hz process. The observed temporal separation of feature encoding extends previous STM studies showing that neural signals during encoding and maintenance can reflect temporally separated memory processing of different scene features (Bocincova & Johnson, 2019; Frenkel & Deouell, n.d.; Martin Cichy et al., 2017). It is also consistent with the behavioral independence of color and orientation reports observed in our data and provides a candidate mechanism for temporally segregating otherwise overlapping feature representations (Bays et al., 2009; Schneegans & Bays, 2017).

The direction of this temporal separation (orientation lagging color during encoding) agrees with behavioral evidence for perceptual asynchrony, in which color is perceived before orientation by approximately 63 ms (Moutoussis & Zeki, 1997; Viviani & Aymoz, 2001). This suggests that features perceived earlier may also be encoded and replayed earlier within the memory-related theta cycle, although this hypothesis requires direct testing. In addition to the main encoding-period analysis, we also examined retrieval-period decoding dynamics (see Appendix). This complementary analysis revealed an inversion of the feature-ordering observed during encoding, with orientation prediction preceding color prediction by approximately 72 ms.

### Spatially distributed theta and beta power contributions

Pairwise comparison of spectral power between correct and error trials revealed stronger frontal and occipital theta power on correct trials during both the encoding and maintenance periods, reproducing established research linking theta to STM performance (Jensen & Tesche, 2002; Sauseng et al., 2010). In contrast, frontal beta power during the encoding-onset window was stronger on trials with poor subsequent memory. This directionality is consistent with reports linking beta desynchronization — that is, beta power decreases — to successful memory formation: Hanslmayr et al. (2014) showed that prefrontal beta power decreased more strongly for subsequently remembered than forgotten items, and that artificially increasing prefrontal beta through rhythmic TMS entrainment selectively impaired encoding, establishing a causal link between elevated prefrontal beta and poorer memory. If reduced beta favors encoding, the elevated frontal beta observed here on poorly remembered trials reflects the converse, less favorable state.

Consistent with these univariate effects, analysis of the MVPA model weights showed that frontal and parietal–occipital channels in the theta and beta ranges contributed most strongly to error prediction. Together, these results indicate that STM fidelity is supported by spatially distributed, multi-frequency activity rather than by a single oscillatory signature or cortical locus (Christophel et al., 2017; Han et al., 2026; Schmidt et al., 2019).

### Attention and saccadic scanning as possible contributors

Theta-rhythmic fluctuations in performance have also been described to underlie attentional processes. Spatial attention can sample multiple locations at ∼4Hz periodicity (Fiebelkorn & Kastner, 2019; Re et al., 2019), with the underlying frontoparietal network alternating between exploitation and exploration states (Fiebelkorn & Kastner, 2019; Sauseng et al., 2010). Under this account, each attentional visit to an item location, e.g. each gaze fixation, would generate a new encoding episode, such that the theta-fluctuation in prediction accuracy would reflect the temporal structure of attentional sampling rather than a memory-specific process.

Several observations argue against this purely attentional explanation. First, error-prediction fluctuations were also present during the maintenance interval, when no stimulus was presented. Second, the fluctuations were not coupled to eye movements: cross-correlation between EOG velocity and decoding time courses revealed no systematic relationship for either feature or load condition (see Appendix). Third, to test whether the same rhythmic architecture operates beyond encoding, we extended the analysis to the retrieval interval (0-3s after spatial-cue onset) and found comparable ∼4 Hz fluctuations, with similar phase offsets across spatial positions and across features, even though the screen showed only a position cue, with no colored or oriented items. Moreover, the temporal relationship between color and orientation reversed relative to encoding During encoding, color prediction lagged behind orientation prediction, whereas during retrieval this ordering was inverted, with color prediction preceding orientation prediction (see Appendix for details). This reversal is consistent with theories proposing that the direction of information flow differs between memory encoding and retrieval (Linde-Domingo et al., 2019; Staresina & Wimber, 2019), although this interpretation remains speculative and requires further investigation.

Nonetheless, attentional sampling and memory processing need not be competing explanations. Both mechanisms fit the embedded-process framework of (Köster & Gruber, 2022), in which theta oscillations serve as a unifying pacemaker for attention and memory and under this view, attentional sampling and memory encoding are not distinct operations (see (Köster & Gruber, 2022).

## Conclusion

Our findings provide direct evidence that working-memory encoding fidelity, indexed by subsequent recall-error prediction, fluctuates rhythmically at theta frequency during encoding and maintenance. These fluctuations were expressed as a distributed, multi-frequency neural pattern that was repeatedly reinstated across the trial. The results further suggest that items and their features are sampled at distinct phases of the theta rhythm, providing a candidate mechanism for segregating overlapping representations. Building on these findings, future work may enable closed-loop approaches that predict memory errors from ongoing neural activity and use temporally targeted stimulation to modulate memory performance.

## Author Contributions

NS: Conceptualization, investigation, data curation, formal analysis, methodology, visualization, writing – original draft, writing – review & editing.

SS: Data curation, writing – original draft.

RR: Data curation, formal analysis, writing – review & editing.

XK: Conceptualization, investigation, methodology, validation, resources, supervision, project administration, data curation, writing – review & editing.

## Ethics Statement

The study was approved by the local ethics committee.

## Resource availability

### Lead contact

Information and requests will be fulfilled by the lead contact, Xenia Kobeleva.

### Materials availability

This study did not generate new unique reagents.

### Data and code availability

Raw de-identified participants’ data have been deposited at Sciebo. They will be publicly available as of the date of publication until.

The original code is available on GitHub (will be publicly available as of the date of publication).

